# Streamlined histone-based fluorescence lifetime imaging microscopy reveals ATM regulation of chromatin compaction

**DOI:** 10.1101/191890

**Authors:** Alice Sherrard, Paul Bishop, Melanie Panagi, Maria Beatriz Villagomez, Dominic Alibhai, Abderrahmane Kaidi

## Abstract

Changes in chromatin compaction are crucial during genomic responses. Thus, methods that enable such measurements are instrumental for investigating genome function. Here, we address this challenge by developing, validating, and streamlining histone-based fluorescence lifetime imaging microscopy (FLIM) that robustly detects chromatin compaction states in fixed and live cells; in 2D and 3D. We present quality-controlled and detailed method that is simpler and faster than previous approches, and uses FLIMfit open-source software. We demonstrate the versatility of our method through its combination with immunofluorescence and its implementation in immortalised cells and primary neurons. Owing to these developments, we applied this method to elucidate the function of the DNA damage response kinase, ATM, in regulating chromatin organisation after genotoxic-stress. We unravelled a role for ATM in regulating chromatin compaction independently of DNA damage. Collectively, we present an adaptable chromatin FLIM method for examining chromatin structure in cells, and establish its broader utility.

## Introduction

In the nucleus, DNA is packaged into chromatin structures (Richmond and Davey, 2003) that determine the activity of genomic DNA in space and time (Dekker et al., 2017; Bickmore, 2013); and may also contribute to non-genetic functions of the genome (Bustin and Misteli, 2016). Such chromatin organisation is underpinned by regulatory epigenetic mechanisms, including histone modifications (Kouzarides, 2007). Visually, interphase chromatin appears to exist in two clearly distinct states: open euchromatin and condensed heterochromatin (Bickmore and van Steensel, 2013). Although these chromatin states seem to be stable at-steady-state conditions, they undergo dynamic reorganisation during genome transduction processes such as transcription (Therizols et al., 2014; Wang et al., 2014), or DNA repair (Lukas et al., 2011; Polo and Almouzni, 2015). Therefore, experimental approaches that enable quantitative analysis of global and regional chromatin compaction states will likely advance our understanding of the principles that govern genome organisation and regulation.

Indeed, advanced cell imaging techniques have proven instrumental in the study of chromatin organisation in intact cells. Cryo-electron tomography (Ou et al., 2017) as well as super-resolution light microscopy methods (Boettiger et al., 2016; Ricci et al., 2015) have provided unprecedented insights into the spatial and temporal organisation of chromatin. In addition, techniques such as fluorescence correlation spectroscopy (FCS) (Toh et al., 2015) and fluorescence anisotropy (Bhattacharya et al., 2009), which involve the use of fluorescently tagged histones (Mora-Bermúdez and Ellenberg, 2007), have allowed analysis of chromatin dynamics. However, a critical limitation of these methods is that they require highly specialised skills and instruments.

To determine chromatin compaction states, Forster (fluorescence) resonance energy transfer (FRET) can be used, where histones are tagged with FRET-compatible fluorescent proteins. FRET measurements can in turn be quantitatively determined by fluorescence lifetime imaging microscopy (FLIM), wherein the lifetime of the donor fluorophore (e.g. GFP) is comparatively measured in the absence or presence of an acceptor probe (e.g. mCherry) (Ishikawa-Ankerhold et al., 2012). This chromatin-FLIM-FRET approach has been previously elegantly performed and validated in live human Hela cells (Llères et al., 2009). However, a key limitation of this previous approach is low scalability as well as difficulties in its applicability, particularly due to the method of generating mammalian cell lines with appropriate distribution of fluorescently tagged histones.

Here, we report the development of a streamlined chromatin FLIM protocol that is simple and scalable. We provide a detailed and adaptable experimental pipeline that allows faster data acquisition, quality-controlled data analysis using open source software, and high experimental reproducibility. As well as validating our chromatin FLIM, we applied this method to shed light on the regulation of chromatin organisation in response to DNA damage.

## Results

In pursuit of developing a simple and robust experimental system, we used lentiviral transduction to derive NIH3T3 cell lines expressing GFP-H2B alone (NIH3T3^GFP-H2B^) or in combination with mCherry-H2B (NIH3T3^H2B-2FP^). We first showed that these cells proliferate at a similar rate to the parental cells (Supplementary Figure 1), confirming that the addition of these fluorophores does not affect cell growth and/or survival. Next, we conducted experiments to measure GFP-H2B fluorescence lifetime in the corresponding cells after paraformaldehyde fixation. The data was analysed using FLIMfit (Warren et al., 2013), an open-source software (see materials and methods for details), which enabled accurate quantification of GFP-H2B fluorescence intensity and fluorescence lifetimes. This software also generates a corresponding fluorescence lifetime (Tau_0_: τ_0_, Figure 1A) map on a pixel-by-pixel basis. For increased accuracy, a merged map is generated wherein each pixel is represented by a colour code, dictated by its fluorescent lifetime value and its brightness determined by the GFP-intensity during FLIM-data acquisition (Figure 1A). This merged map is highly informative as it depicts the fluorescence lifetime (Tau, τ), as well as GFP-intensity, and enables the visualisation of a spectrum of chromatin compaction states within the nucleus (Figure 1A). Importantly, FLIMfit software also provides graphical representation of chi-squared (χ^2^) values, a statistical test that indicates the extent of variation between the actual FLIM measurements and the data-fitting model, thus providing a quality-control for validating FLIM data analyses (Figure 1A). To this end, we found that the mean GFP-H2B fluorescence lifetime was discernibly higher in NIH3T3^GFP-H2B^, compared to NIH3T3^H2B-2FP^, at the single nucleus level (Figure 1A). This difference can also be observed as a shift in the distribution of GFP-H2B fluorescence lifetime values of each pixel (Figure 1B); which is consistent with specific FRET from GFP-H2B (donor) to mCherry-H2B (acceptor), when they are co-expressed in NIH3T3^H^2^B-2FP^ cells. Importantly, we could also extend the utility of the chromatin FLIM approach to human retinal pigment epithelial-1 (RPE1) cells (Supplementary Figure 2A and B). Therefore, our findings establish a versatile experimental approach for performing FLIM in fixed cells, to detect FRET between two histones.

**Figure 1.**
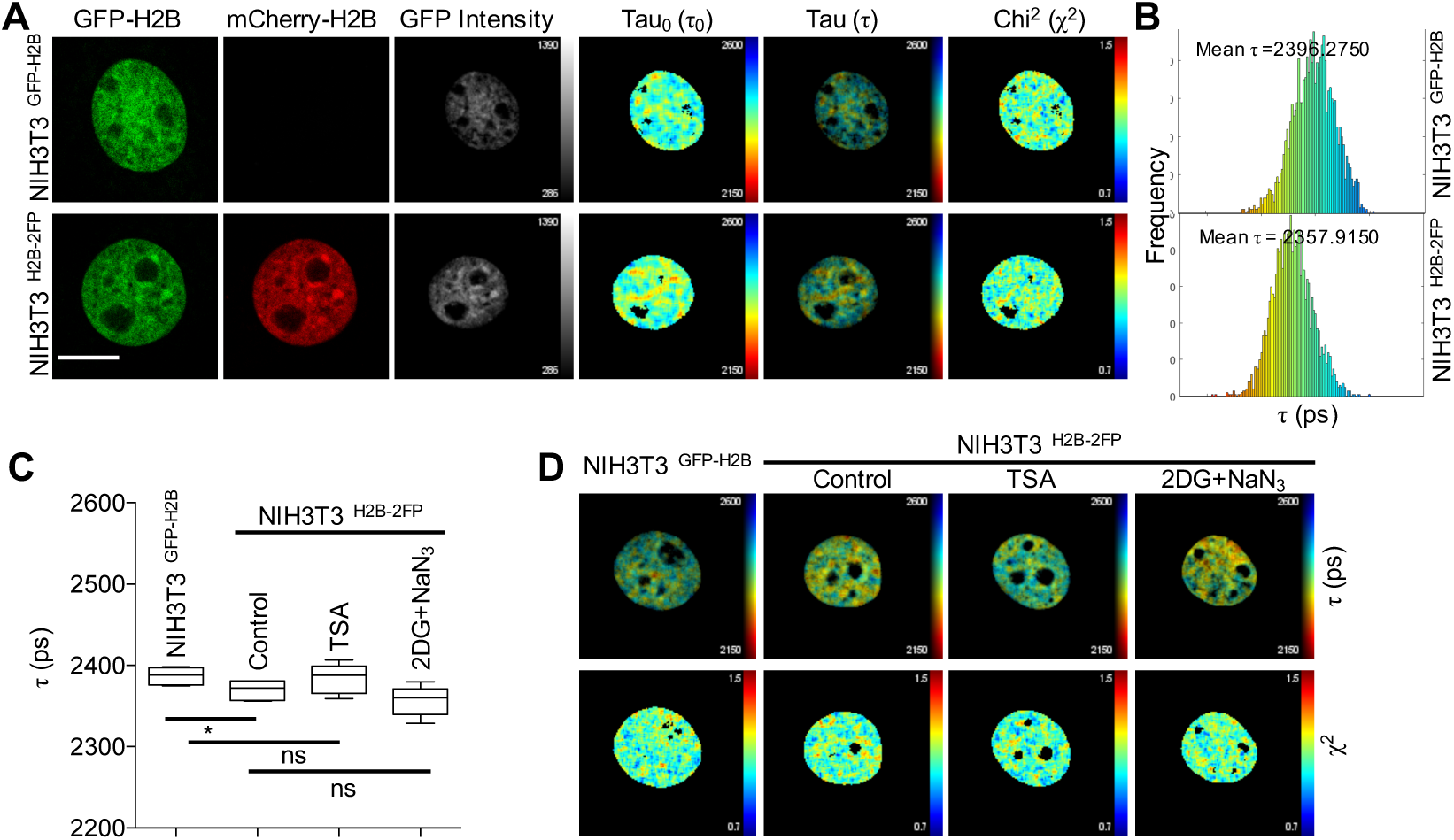
Establishment of chromatin FLIM assay in NIH3T3 cells. **A.** Examples of the FLIM measurement pipeline in the indicated NIH3T3 cells. Images for GFP-H2B and mCherry-H2B were acquired (scale bar = 10 μm), and FLIM data analysis was performed on FLIMfit software that generates the corresponding: GFP intensity, GFP fluorescence lifetime Tau_0_ (τ_0_), merged fluorescence lifetime Tau (τ), chi-squared (χ^2^) maps (please note that the raw images have been re-sized for publication), and **B** the distribution of Tau (τ) values on a pixel-by-pixel basis of the same nuclei shown in a. **C.** Quantification of τ in the indicated cells and growth conditions. Data expressed in picoseconds (ps) as Mean τ±SD, n=5, * indicates *p*=0.0422, ns indicates *not-significant*, Student’s *t*-test. **D.** Example fluorescence lifetime Tau (τ), chi-squared (χ^2^) maps from **C**.

A key, yet currently unmet, challenge in performing chromatin FLIM in mammalian cells is the ability to conduct FLIM measurements in large numbers of mammalian cells (Llères et al., 2009). Addressing this limitation would not only provide experimental robustness through statistical power, but would also allow the identification of potential heterogeneity within a population of cells. Accordingly, we analysed GFP-H2B fluorescence lifetime in multiple fixed NIH3T3^GFP-H2B^ and NIH3T3^H2B-2FP^ cells from the same experiment. As shown (Figure 1C), GFP-H2B fluorescence lifetime in NIH3T3^GFP-H2B^ was significantly higher than in NIH3T3^H2B-2FP^, thus confirming our single cell analysis.

Changes in GFP-H2B fluorescence lifetime in NIH3T3^H2B-2FP^ cells can be used to infer the proximity between GFP-H2B and mCherry-H2B (typically in the range of ∼1-10nm), and therefore it relates to the extent of chromatin compaction (Llères et al., 2009). When chromatin is in its de-compact state, nucleosomes are spaced apart, leading to higher fluorescence lifetime of the donor fluorophore (due to a decrease in FRET from to the donor to acceptor fluorophores (Ishikawa-Ankerhold et al., 2012)). Whereas when chromatin is in its compact state, nucleosomes are closer to one another, leading to an increase in FRET and a decrease in the donor fluorescence lifetime. As such, we used growth conditions to alter chromatin compaction in interphase NIH3T3^H2B-2FP^ cells. Here we found that inducing chromatin relaxation using the HDAC inhibitor Trichostatin A (TSA) did not significantly affect GFP-H2B fluorescence lifetime at a cell population level (Figure 1C). Similarly, increasing chromatin compaction using 2-deoxy-glucose (2DG) and sodium-azide (NaN_3_)-mediated ATP depletion had no significant effect on GFP-H2B fluorescence lifetime in a cell population (Figure 1C). However, when examining GFP-H2B fluorescence lifetime at the single cell level, we could detect an increase (with TSA treatment) or decrease (with 2DG+NaN_3_ treatment), consistent with previous findings (Examples in Figure 1D).

Given that both TSA and 2DG+NaN_3_ treatments are well documented to alter of chromatin condensation state, our observed heterogeneity in GFP-H2B fluorescence lifetime between cells could be attributed to the relative levels of GFP-H2B and mCherry-H2B expression in NIH3T3^H2B-2FP^ cells. To test this, we reasoned to perform fluorescence activated cell sorting (FACS) to generate different NIH3T3^H2B-2FP^ cell subpopulations with varying expression levels of GFP-H2B and mCherry-H2B. We sorted eight different subpopulations of NIH3T3^H2B-2FP^ (Supplementary Figure 3A) and performed FLIM experiments to determine the extent of heterogeneity in GFP-H2B fluorescence lifetime. We observed that subpopulations 3 and 6 had the lowest degree of variation in GFP-H2B fluorescence lifetime (Supplementary Figure 3B). To minimise any effects of extreme protein overexpression (in subpopulation 3), we used subpopulation 6 for the subsequent experiments. By using this subpopulation in FLIM experiments, we found that TSA treatment resulted in a significant increase in GFP-H2B fluorescence lifetime at a population level and that 2DG+NaN_3_ decreased GFP-H2B fluorescence lifetime (Figure 2A-B). Therefore, by controlling the expression of GFP-H2B and mCherry-H2B, we established a robust method that allows quantitative analysis of chromatin compaction states in fixed cells, and importantly at a population level.

**Fig. 2.**
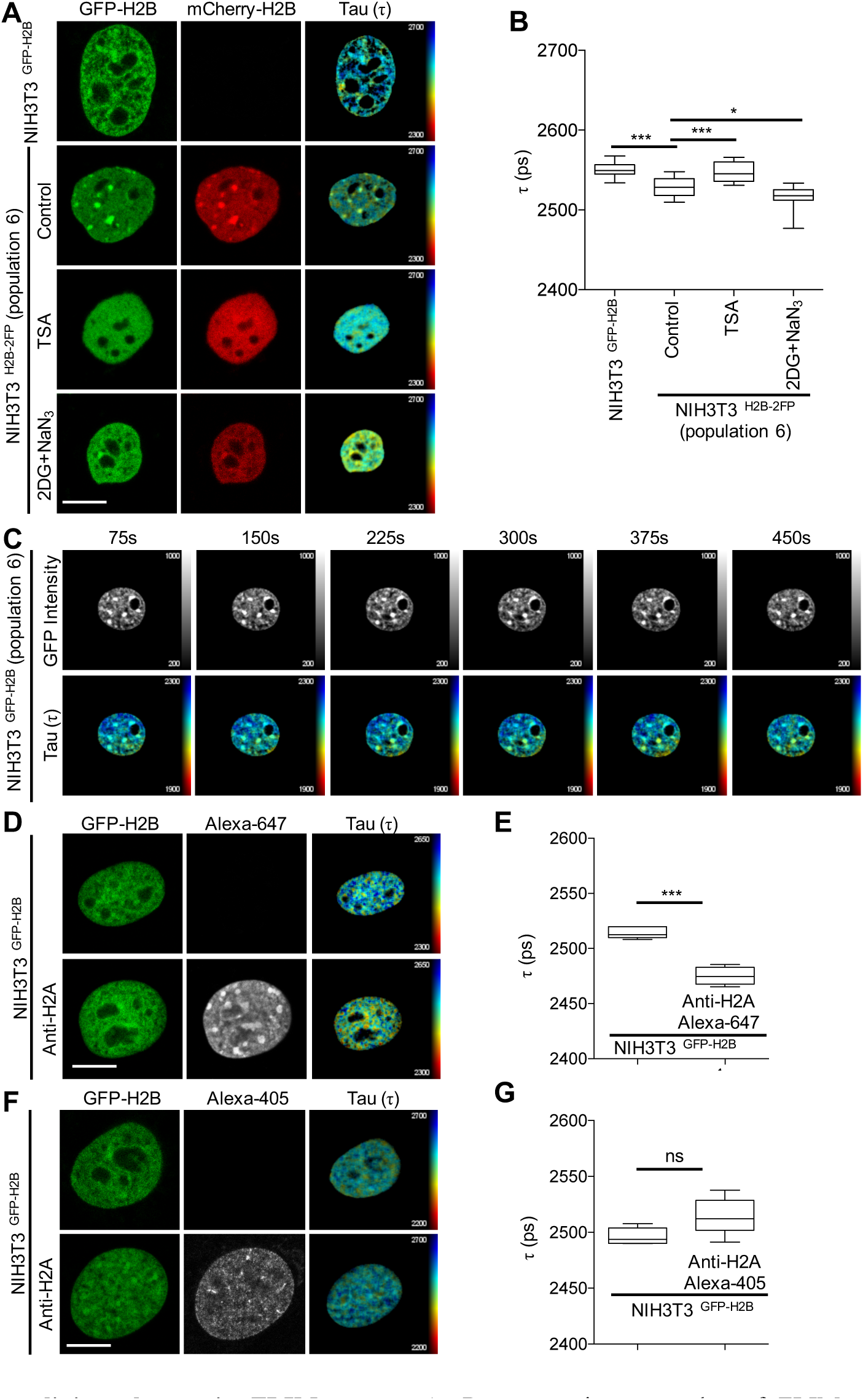
Streamlining chromatin FLIM assays. **A.** Representative examples of FLIM measurements conducted in the indicated cells and conditions (scale bar = 10 μm). **B.** Quantifications of τ data from **A** expressed as Mean τ±SD, n≧13, * indicates *p*=0.0312, *** indicates p≦0.001, Student’s *t*-test. **C.** Live-cell chromatin FLIM in NIH3T3^H2B-2FP^ at the indicated time intervals. **D.** Example of GFP-H2B τ in fixed NIH3T3^H2B-2FP^ co-stained with Anti-H2A using AlexaFluor-647 (scale bar = 10 μm). **E.** Quantification of τ data from **D** expressed as Mean τ±SD, n≧5, *** indicates p≦0.001, Student’s *t*-test. **F.** Similar to **D** but using AlexaFluor-405. **G.** Quantification of **F**, n≧5, ns indicates *not-significant*, Student’s *t*-test.

Noteworthy, it did not escape our attention that this method of cell line generation, through the use of FACS, is not suitable for all cell types. Therefore, to increase the versatility of our method, we applied an alternative approach using cultured primary neurons (due to their incompatibility with FACS) as an example. Here, we used pre-extraction with cytoskeleton buffer (CSK) (Kaidi et al., 2010) prior to cell fixation to remove free non-incorporated fluorescently tagged histones, thus minimising variation in the levels of GFP-H2B and mCherry-H2B proteins; which we identified earlier to be a critical factor in the feasibility of our FLIM approach. Accordingly, pre-extraction allowed us to measure fluorescence lifetime changes as well as chromatin relaxation induced by TSA, in both unsorted NIH-3T3 cells (Supplementary Figure 4A-B), as well as in primary neurons (Supplementary Figure 4C-D).

Upon the development of this experimental system, we next optimised FLIM data acquisition, to increase the speed at which FLIM measurements can be taken. By adjusting acquisition parameters (see materials and methods), we were able to conduct chromatin FLIM in live cells at time intervals of 75 seconds (the fastest rate of chromatin FLIM measurements in mammalian cells to date (Llères et al., 2009)), providing scope for rapid chromatin rearrangements to be detected in real time (Figure 2C). Collectively, our method extends on the previous elegant work that allowed chromatin-based FLIM quantification in live cells (Llères et al., 2009; 2017).

Another advantage of our FLIM approach in paraformaldehyde fixed cells, is the possibility of combining it with immunofluorescence staining. Indeed, this could be useful when investigating changes to chromatin organisation, for example within specific nuclear compartments or in response to cellular stimuli. However, a key consideration would be to ensure that fluorescent antibodies do not interfere with chromatin FLIM. Accordingly, we stained for histone H2A in FACS-sorted NIH3T3 ^GFP-H2B^ cells using either AlexaFluor-405 or AlexaFluor-647 secondary antibodies, followed by measuring GFP-H2B fluorescence lifetime. The results revealed that the presence of AlexaFluor-647 significantly interferes with GFP-H2B fluorescence lifetime (Figure 2D-E), whereas using AlexFluor-405 had no discernible effect (Figure 2F-G). This important control suggests that parallel AlexaFluor-405 staining is compatible with performing FLIM experiments using the GFP-mCherry FRET pair.

Having established a robust chromatin FLIM approach that can be combined with conventional immunocytochemistry, we applied this method to determine the regulation of chromatin organisation in response to DNA damage. We treated NIH3T3^H2B-2FP^ cells with the topoisomerase II inhibitor etoposide (ETP) and confirmed DNA damage induction by the presence of H2A.X-phospho-S139 (γH2AX) and KAP1-phospho-S824 (pKAP1) (Figure 3A and Supplementary Figure 5A). When analysing DNA-damaged γH2AX positive cells, we found that DNA damage resulted in increased GFP-H2B fluorescence lifetime (Figure 3A-B), suggestive of increased chromatin relaxation (Takahashi and Kaneko, 1985). Notably, inhibition of the DNA damage kinase ATM abrogated etoposide-induced chromatin de-compaction (Figure 3A-B and Supplementary Figure 5A), consistent with previous reports of ATM-dependent chromatin relaxation after DNA damage (Ziv et al., 2006; Caron et al., 2015). This data highlights the usefulness of our chromatin FLIM method in revealing the structural responsiveness of chromatin to genotoxic stress.

**Figure 3.**
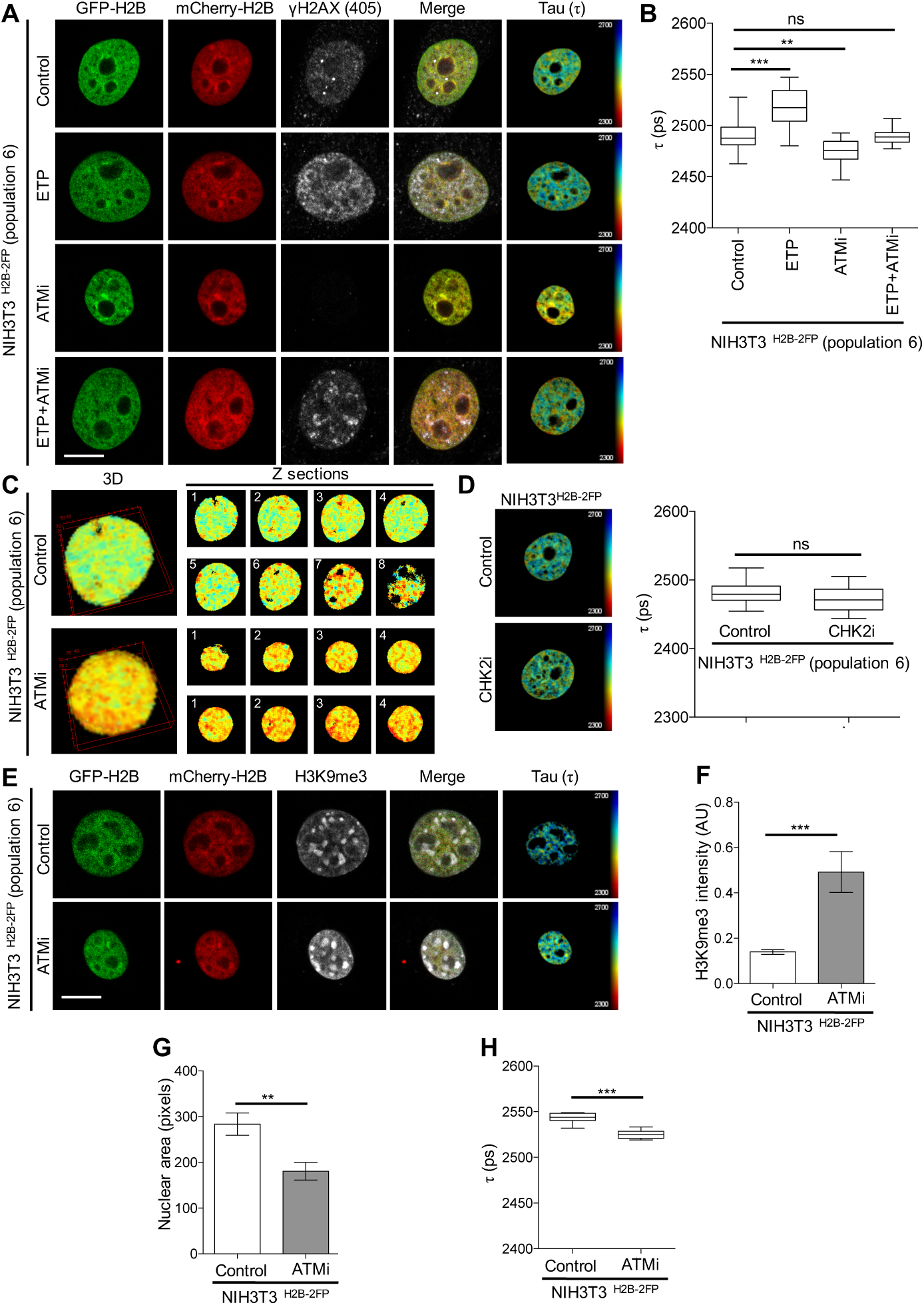
ATM regulates chromatin compaction state. **A.** Representative examples of FLIM measurements conducted in the indicated cells that were treated as indicated and co-stained for γH2AX (scale bar = 10 μm). **B.** Quantifications of τ data from **A** expressed as Mean τ±SD, n≧18, ** indicates *p*=0.010, *** indicates *p*≦0.001, Student’s *t*-test. **C.** Example of 3D-FLIM in NIH3T3^H2B-2FP^ cells treated with ATMi; the right panels are different Z sections from **C** (with lower zoom). **D.** *Left panel*: Example of GFP-H2B τ in fixed NIH3T3^H2B-2FP^ treated with CHK2i; *right panel*: corresponding quantifications, n≧16, *ns* indicates *not-significant* p=0.075, Student’s *t*-test. **E.** Simultaneous analyses of τ and intensity of H3K9me3 in the same NIH3T3^H2B-2FP^ treated with ATMi, and their corresponding quantification (n=10): H3K9me3 (**F**, *** indicates *p*≦0.001), nuclear area (**G**, ** indicates *p*=0.004), and τ (**H**, *** indicates *p*≦0.001); Student’s *t*-test.

Notably, we observed that ATM inhibition alone resulted in a marked decreased in GFP-H2B fluorescence lifetime in 2D (Figure 3A-B). Similarly, we could detect decreased GFP-H2B fluorescence lifetime (i.e. increased chromatin compaction) in 3D (Figure 3C). Furthermore, inhibition of CHK2, a downstream kinase in ATM-dependent signalling (Matsuoka et al., 2000), did not result in increased chromatin compaction, suggesting that ATM is not signalling through its canonical pathway (Figure 3D and Supplementary Figure 5B). These observations suggest that basal ATM activity may be somewhat required for maintaining chromatin organisation in interphase cells, in a manner that involves the regulation of condensed chromatin (heterochromatin). To further investigate this, we performed FLIM measurements and analysed the heterochromatic histone mark H3K9me3 (histone H3 lysine-9 tri-methylation) (Bannister et al., 2001) in the same cells, while also measuring nuclear area. These concurrent analyses revealed that ATM inhibition resulted in a marked increase in H3K9me3 fluorescence intensity (Figure 3E-F), and a significant decrease in nuclear area (Figure 3E-G) as well as reduced fluorescence lifetime (Figure 3E-H). In light of these findings, we conducted a reltively higher throughput imaging screen in ATM inhibited cells in 3D to examine proteins and modifications that are associated with heterochromatin (Kouzarides, 2007). Results showed an increase in specific heterochromatin markers including H3K9me3 and HP1γ and H3K27me3 (Supplementary Figure 5C-E), but not in H3K9me2 and HP1α (Supplementary Figure 5E-F). Collectively, these findings reveal a hitherto unappreciated role for ATM basal kinase activity in establishing chromatin organisation through the regulation of specific heterochromatin factors.

## Discussion

Here we have developed an adaptable and transferable method, using fluorescence lifetime imaging microscopy to detect chromatin compaction states in live and fixed cells both in 2D and 3D. Unlike a previous method, we use pulsed-laser (not multiphoton confocal) and open source fitting software, FLIMfit for data analysis. Thus, our chromatin-FLIM method offers a wider scope for its utility and applicability. Notably, chromatin FLIM data analysis through FLIMfit provides chi-squared (χ^2^) maps as a quality control step for suitability of the FLIM data-fitting model (Warren et al., 2013). These advances provide a streamlined protocol to conduct FLIM experiments in a large number of cells and in different cell types (mouse fibroblasts, human epithelial cells, and primary rat neurons). These advantages allow for the identification of population-based heterogeneity and increased statistical power. Furthermore, our method is amenable to modification and can be adapted to automation for the purpose of screening. This may be useful to identify agents and/or factors that influence chromatin compaction states, with potential to revealing the mechanisms that underlie genome organisation at a system level. This is particularly relevant given that FLIMfit is part of the Open Microscopy Environment (OME) (Jason, 2013) that seamlessly integrates large data sets from screens. Relevant to this, through the use of pixel-based segmentation and combined antibody labelling, our FLIM method could give insights into the association of condensed/de-condensed chromatin with sub-nuclear domains, or chromosome territories, thus increasing our understanding of genome compartmentalisation (Bickmore and van Steensel, 2013). Furthermore, we have increased the speed at which such measurements can be taken, thus providing a system that is compatible with time-dependent processes, and the potential identification of previously unresolved alterations to chromatin compaction in response to stimuli. Noteworthy, such advances have recently been made in Caenorhabditis elegans (C.elegans) (Llères et al., 2017), and our method extends them to mammalian cells.

Highlighting its utility, we applied our method to provide the first evidence for global chromatin relaxation upon DNA damage in intact cells. We further show that this chromatin relaxation upon DNA damage is ATM-dependent, which is particularly pertinent given the role of ATM in promoting DNA repair within heterochromatin (Goodarzi et al., 2008). Of particular interest, we also observed that inhibition of ATM in resting cells resulted in pronounced global chromatin compaction, a finding which correlated with increased H3K9me3 and reduced nuclear volume. This suggests a role of ATM in chromatin surveillance, likely outside of a DNA damage context, and is consistent with previous findings (Bakkenist and Kastan, 2003; Kaidi and Jackson, 2013). It is possible that the reported ATM-dependent transcriptional inhibition after DNA damage (Shanbhag et al., 2010; Kruhlak et al., 2007) is somewhat mediated through the ability of ATM to regulate chromatin compaction state. The prospect of ATM kinase as a key regulator of chromatin organisation, beyond its role in the DNA damage response, is an exciting one. Indeed, this ATM function may help to explain its role in neurodegeneration (Jackson and Bartek, 2009) wherein heterochromatin appears to be dysregulated (Frost et al., 2014). Given its homology with ATR (Awasthi et al., 2016), which has been reported to signal during cellular mechanotransduction (Kumar et al., 2014), it is conceivable to speculate that ATM may regulate chromatin compaction states during mechanical cell signalling events that are known to alter chromatin organisation (Shivashankar, 2011).

In summary, we have developed and validated a versatile method to assess chromatin compaction states in intact cells, and applied this method to provide novel insights into the regulation of chromatin compaction by ATM. Given the importance of chromatin compaction states in regulating genome transduction processes such as transcription and DNA repair, we envisage that future research may take advantage of our chromatin FLIM method to further the mechanistic understanding of spatiotemporal control of genome organisation and function, both in health and disease.

## Materials and Methods

### Cell culture and treatments

NIH3T3 cells were cultured in in Dulbecco’s Modified Eagle’s Medium (DMEM) and RPE1 cells in DMEM-F12 Ham (Gibco), containing 2 mM glutamine (Sigma), 100 units/ml Penicillin (Gibco), 100 g/ml Streptomysin (Gibco) and 10% foetal bovine serum (FBS) (Gibco). Cells are maintained at 37°C in 5% CO_2_ dry incubators. For FLIM experiments, cells were seeded in glass μ-dishes (ibidi). Cells were treated with 1 μM TSA (Sigma) for 16 hrs, 50 mM 2-Deoxyglucose (Sigma) and 10 mM Sodium Azide (Sigma) for 30 minutes, 10 μM ATM inhibitor ATMi (Abcam, KU-55933), or 1.5 μM CHK2i (Sigma, C3742) both for 18 h before treatment 10 μM etoposide (Sigma) for 1 h.

Primary hippocampal neuronal cultures were prepared from embryonic day E18 rats and maintained using standard conditions. Neurons were grown on 35 mm glass-bottom dishes (ibidi). Day-*in-vitro* 2 (DIV2) neurons were infected with GFP-H2B and mCherry-H2B lentiviruses. On DIV6, neurons were treated with 55 mM KCl for 3 minutes to increase histone exchange. On DIV7, neurons were then pre-extracted and fixed as described. Animal care and all experimental procedures were conducted in accordance with UK Home Office and University of Bristol guidelines.

### Antibodies

Antibodies were obtained from Cell Signalling (H2A: 12349P; H3K27me3: 9733S; DyLight-680: 5470 and Dylight-800: 5151), Abcam (AlexaFluor-405: ab175651; H3K9me3: ab8898; H2A.X: ab124781; KAP1: ab10483; KAP1-pS824: ab70369; HP1γ: ab10480 and H3K9me2: ab1220), and Invitrogen (AlexaFluor-647, A21245) and Millipore (γH2A.X, 05-636; HP1α: 05-689).

### Immunofluorescence staining

Cells were fixed in 4% paraformaldehyde for 10 minutes, permeabilised in 0.1% Triton X-100 for 10 minutes, and blocked in 1% BSA in PBS for 30 minutes. For pre-extraction experiments, cells were incubated in CSK buffer (10 mM PIPES, pH 6.8, 100 mM NaCl, 300 mM Sucrose, 3 mM MgCl_2_, 1 mM EGTA, 0.5% Triton X-100) for 5 minutes before fixation. Cells were incubated with primary antibody for 2 h, and secondary antibody for 45 minutes, both at RT.

### Production of lentivirus

To produce lentivirus, the *PGK-GFP-H2B* and *PGK-H2B-mCherry* vectors were obtained from Addgene. Each of these plasmids was co-transfected with helper plasmids into HEK293T packaging cells. Lentivirus particles were recovered using standard protocols.

### Fluorescence-activated cell sorting (FACS)

GFP-H2B and mCherry-H2B expressing NIH3T3 cells were isolated using fluorescence-activated cell sorting (FACS), and sorted into eight populations (see FACS plots). Viable cells were identified based on light scatter and the exclusion of propidium iodide (PI). In addition, single cell gating was used to exclude doublets and aggregated cells. GFP-H2B and mCherry-H2B expressing cells were sorted using 488nm laser excitation and 510-550nm emission and 552nm excitation with 600-620nm emission, respectively, using a Becton Dickinson InFlux cell sorter (BD Biosciences, Franklin Lakes, NJ) running BD Software version 1.2.

### Fluorescence lifetime imaging microscopy (FLIM) data acquisition

FLIM was performed on cells growing on glass μ-dishes (ibidi). In the case of fixed samples, no mounting medium was used, and FLIM was performed in PBS. Fluorescence lifetime images were acquired on a Leica TCS SP8 system attached to a Leica DMi8 inverted microscope (Leica Microsystems). Excitation was provided by a white light laser with a repetition rate of 20 MHz and an acousto-optical beam splitter (AOBS) selected an excitation wavelength of 488 nm. Excitation continued for 75 seconds per FLIM measurement/focal plane. Images were acquired using a 63x 1.4 NA oil immersion objective. Fluorescence of the GFP-H2B was detected using a hybrid detector operating in photon counting mode over an emission range of 495 – 530 nm. A notch filter centred on 488nm minimised any laser scatter into the detector. Time-resolved data was acquired through use of a PicoHarp 300 TCSPC module (PicoQuant) controlled through SymPhoTime64 software (PicoQuant). FLIM Images were acquired with 512 x 512 pixels and 4096 time bins. For live cell experiments, the system was maintained at 37°C/5% CO2. In 3D experiments, FLIM measurements were taken at a minimum of 8 focal planes, 1μm apart.

### FLIM data fitting

Fitting of FLIM images was performed with the FLIMfit software tool (version 5.0.3) developed at Imperial College London. Temporal binning of the fluorescence decays was performed prior to fitting, resulting in 401 time bins per decay and the images were spatially binned 4x4 to ensure sufficient photons were present per pixel prior to the fitting of the data. Fitting of the fluorescence images was then performed pixelwise with a single exponential model on all pixels above an intensity threshold of 200 photons with a 5x5 smoothing kernel applied, allowing spatial variations in fluorescence lifetime to be visualised. The instrument response function (IRF) was measured by imaging a solution of 1 micromolar rhodamine 6G with the same settings as data acquisition and then using the “Estimate IRF” function within the FLIMfit software to extract the IRF.

### Imaging data analysis

Fiji software was used to segment the nucleus, based on the GFP-H2B channel. The size of the segmented nuclear area (in pixels), and the intensity of H3K9me3 within it were then quantified using Fiji software.

### High throughput quantitative imaging

Imaging was performed on a Perkin Elmer Opera LX system, using a 20x water objective, imaging in 3D at 1 μm per Z-plane. Image analyses of maximum-intensity-projections were conducted using a customised MATLAB pipeline.

### Statistics

Statistical analysis was performed on GraphPad Prism, using Student’s *t-*test to compare selected pairs, as indicated.

## Author contribution

AK conceived and supervised the project. AS conducted most experiments with significant contributions from: PB (neuronal experiments, and FLIMfit), MP (immunoblotting and immunofluorescence), DA (FLIM experimental optimisations, and FLIMfit). MBV generated lentiviruses. AK and AS wrote the manuscript with contributions from all authors.

## Acknowledgments

We wish to acknowledge the assistance of Dr Andrew Herman and Lorena Sueiro Ballesteros for cell sorting and the Faculty of Biomedical Sciences Flow Cytometry Facility, University of Bristol. We acknowledge the Wolfson Bioimaging Facility and are specifically grateful to Dr Stephen Cross for assistance with imaging data analysis. AK laboratory is funded by MRC New Investigator Award (MR/N000013/1), HFSP Program Grant (RGP0021/2016) with infrastructure from a Wellcome Trust Seed Awards in Science (WT107789AIA). AS is funded by University of Bristol PhD scholarship. MP is funded by a Wellcome Trust Dynamic Cell Biology PhD scholarship. FLIM was carried out the at the Wolfson Bioimaging Facility, University of Bristol, through BrisSynBio, a BBSRC/EPSRC-funded Synthetic Biology Research Centre (grant number: L01386X)

---

**Supplementary Figure 1.**
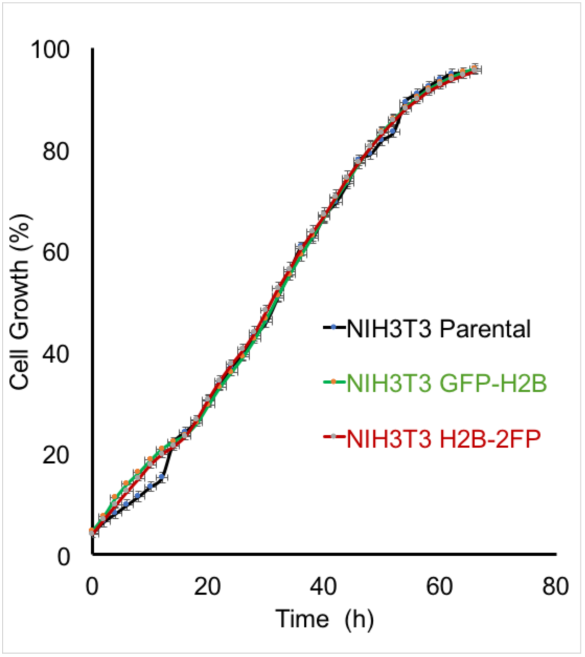
Analysis of the growth rate of NIH3T3. The proliferation rates of the indicated NIH3T3 cell derivatives were quantitatively analysed using an automated live cell imaging system (Incucyte-ZOOM), data from one experiment performed in triplicates are presented as Mean±SE.

**Supplementary Figure 2.**
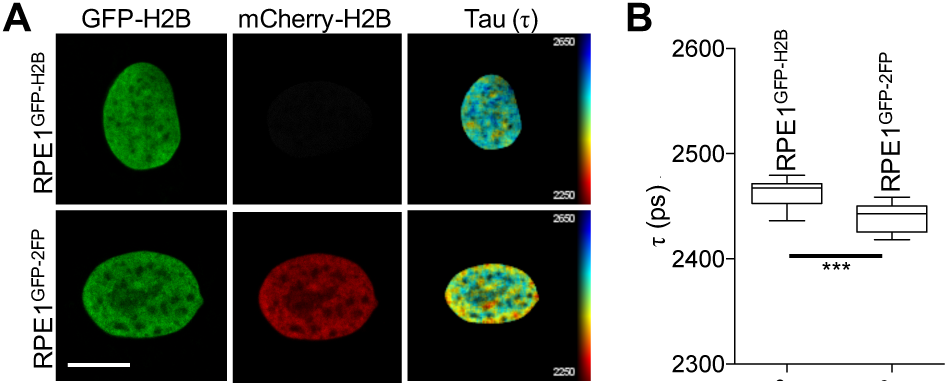
Establishment of chromatin FLIM assay in RPE1 cells. **A.** Examples of FLIM measurements in human RPE1 cells as indicated (H2B images’ scale bar = 10 μm). **B.** Quantification of τ in multiple RPE1 cells as in **A**. Data expressed in picoseconds (ps) as Mean τ±SD, n=10, *** indicates *p* = 0.0012, Student’s *t*-test.

**Supplementary Figure 3.**
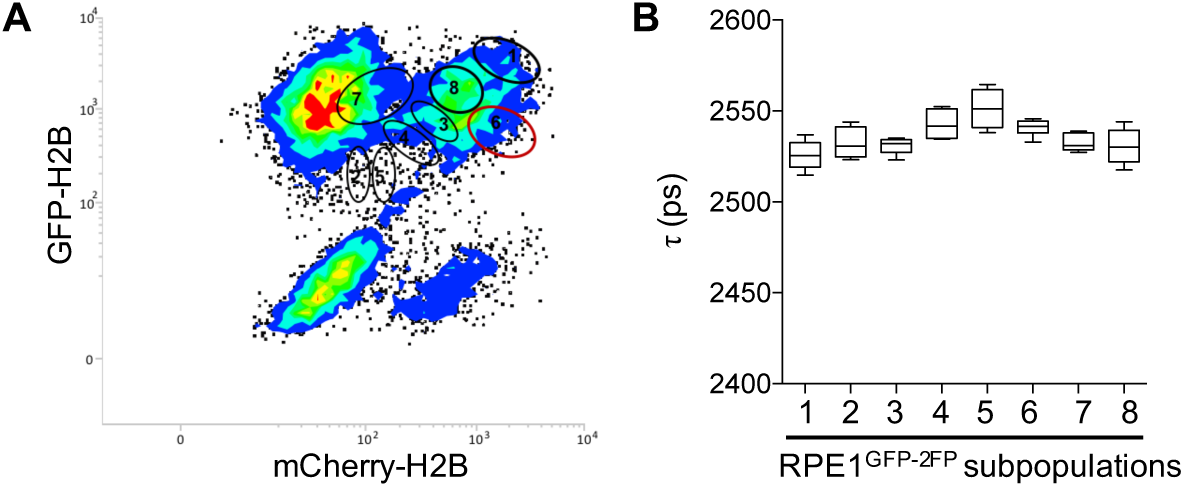
Optimising the expression levels of fluorescently-tagged histones for chromatin FLIM using FACS. **A.** Two-dimensional scatter plot for NIH3T3 cells co-expressing GFP-H2B and mCherry-H2B (NIH3T3^H2B-2FP^). The indicated numbers (1-8) correspond to FACS-sorted subpopulations of cells that express varying levels of GFP-H2B and mCherry-H2B. **B.** Quantification of τ in multiple FACS-sorted subpopulations of NIH3T3^H2B-2FP^ cells from **A**, and data are expressed in picoseconds (ps) as Mean τ ± SD, n=5.

**Supplementary Figure 3.**
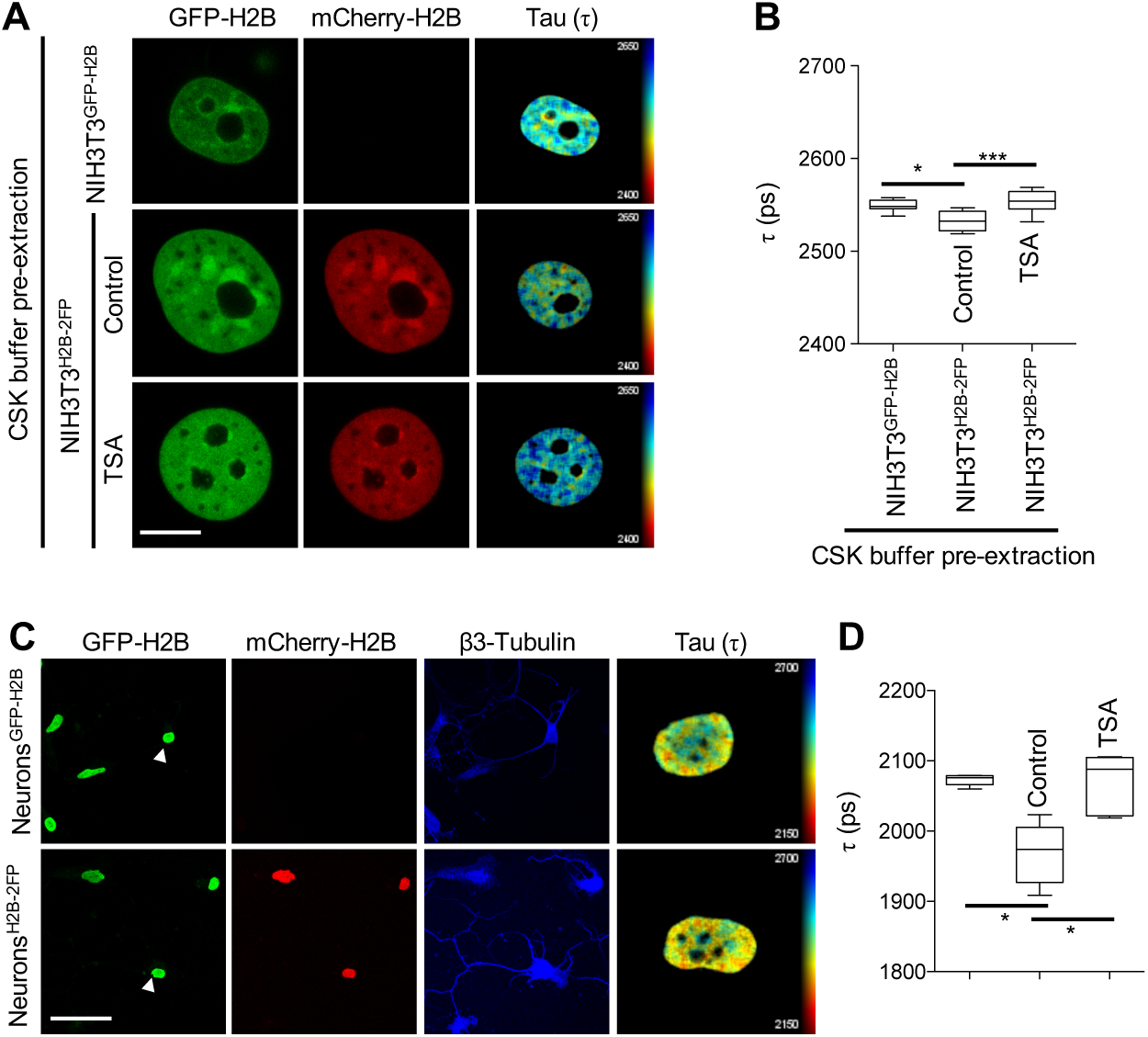
Optimising the levels of fluorescently-tagged histones for chromatin FLIM using pre-extraction. **A.** Representative examples of FLIM measurements conducted in the indicated cells and conditions after CSK-buffer pre-extraction and fixation (scale bar = 10 μm). **B.** Quantification of τ data from **A**, expressed in picoseconds (ps) as Mean τ±SD, n≧7, * indicates *p*=0.03, *** indicates p<0.001, Student’s *t*-test. **C.** Representative examples of FLIM measurements conducted in hippocampal neurons after CSK-buffer pre-extraction and fixation (scale bar = 100 μm, the FLIM analysed nuclei are indicated by white arrows). **D.** Quantification of τ data from **C**, expressed in picoseconds (ps) as Mean τ±SD, n≧5, * indicates *p*=0.03, Student’s *t*-test.

**Supplementary Figure 5.**
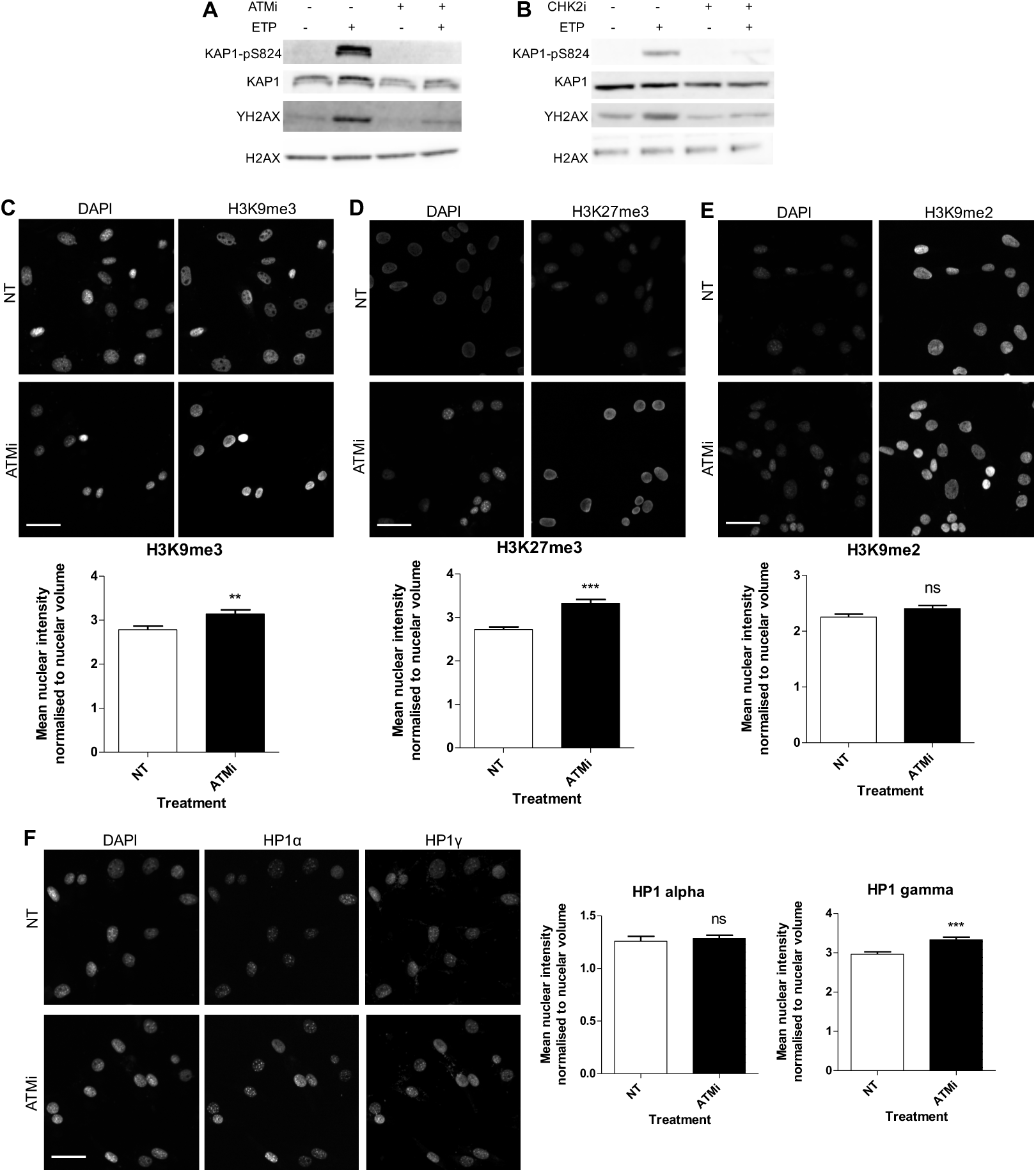
ATM regulates heterochromatin-associated histone modifications. **A and B.** Immuno-blot analyses of NIH3T3 treated as indicated. **C** to **F.** Representative images (scale bar = 50 μm) of the indicated histone modification and their corresponding quantifications in NIH3T3 treated with ATMi, presented as Mean±SD, n≧750, *** indicates *p*<0.001, ** indicates *p*=0.005; Student’s *t*-test.

